# Patterns and correlates in the distribution, design and management of garden ponds along an urban-rural gradient

**DOI:** 10.1101/2023.11.30.569344

**Authors:** Andrew J. Hamer, Barbara Barta, Zsuzsanna Márton, Csaba F. Vad, Beáta Szabó, Irene Tornero, Zsófia Horváth

## Abstract

Urbanisation results in the loss and alteration of natural wetlands and ponds. However, garden ponds in cities and towns can act as rich reservoirs of aquatic biodiversity and stepping stones for dispersal. Homeowners with a range of different motivations, including biodiversity values, install garden ponds. Here, our main aim was to study whether the design and management choices of garden pond owners was dependent on the location of ponds (capital city vs. countryside), when ponds were installed (pond age), or whether fish were introduced. We surveyed 834 garden pond owners across Hungary using a citizen science questionnaire, asking questions on pond size, location, construction date and materials, vegetation structure, introduction of fish and management practices. From 753 validated responses, we found that the introduction of fish into ponds and high urbanisation were strongly associated with local features and management practices, especially large ponds with a water circulation feature, irrespective of pond age. A typical garden pond in Hungary is ∼20 m^2^, <10 years old, made of rubber (PVC) lining, contains fish, aquatic vegetation and circulating water, and is actively managed. There was a spatial separation of ponds based on local features between ponds in the capital city (Budapest) and elsewhere. These findings suggest that garden pond owners in cities are likely to make different choices in pond design and management compared to owners in regional areas. Our results also suggest that pond owners may primarily select management practices to improve habitat quality for ornamental fish. Our findings have important implications for maintaining aquatic biodiversity in urban areas, where garden ponds may be the only aquatic habitat available.

**RESEARCH HIGHLIGHTS:**

- Hungarian garden ponds ≤ 200 m^2^ occur in both large cities and small villages
- Most garden ponds we sampled in Hungary were built in the last decade
- Pond design and management differs between Budapest and regional areas
- Pond design and management practices are influenced by ornamental fish stocking
- Pond owners frequently drain ponds to remove sediment and leaves and use algaecides

## 1. INTRODUCTION

Urbanisation is occurring rapidly in parallel with global human population growth, leading to the loss and modification of natural habitats (McKinney, 2002). Between 2000 and 2030, the global urban population is forecast to reach nearly 5 billion, and urban land cover will increase by 1.2 million km^2^ (Seto et al., 2012), with 290,000 km^2^ of natural habitat converted to urban land uses (McDonald et al., 2020). Freshwater habitats are a valuable natural resource but are one of the most threatened ecosystems (Sala et al., 2000; Zedler and Kercher, 2005), largely due to land-use change and human activities (Dudgeon et al., 2006; Reid et al., 2019). Urbanisation has contributed greatly to the loss and degradation of freshwater ponds and wetlands (Burgin et al., 2016; Ehrenfeld, 2000; Kentula et al., 2004), with high rates of pond loss being particularly evident in Europe (Biggs et al., 2005; Smith et al., 2022; Wood et al., 2003), resulting in increased pond isolation (Thornhill et al., 2018). Small ponds and pond networks provide crucial ecosystem services – improving human well-being by providing aesthetic enjoyment and leisure, providing habitats for biodiversity, acting as carbon sinks, and mitigating climate change (Cuenca-Cambronero et al., 2023; Oertli et al., 2005). Yet, despite profound losses in this vital resource since the twentieth century, there are opportunities to increase the area of freshwater habitat in urban areas. For example, garden ponds may provide one solution to the loss of ponds in urban and agricultural areas, potentially offsetting losses of natural ponds (Gledhill et al., 2008).

Garden ponds are small artificial waterbodies, typically in the size range of other lentic freshwater habitats (1 m^2^ – 2 ha; Biggs et al., 1994; Hill et al., 2021), built within private gardens and set within a broader matrix of landscape imperviousness (Hassall, 2014). Homeowners build or install garden ponds to improve the aesthetic value of gardens but also to support ornamental fish and biodiversity (Blicharska and Johansson, 2016; Hill et al., 2021). Due to their popularity, garden ponds can contribute substantially to the overall cover of standing freshwater. For example, in the United Kingdom (UK) there are an estimated 2.5 – 3.5 million garden ponds equating to 349 ha of standing water, although the total area is fragmented into tiny patches and distributed over a wide area (Davies et al., 2009; Gaston et al., 2005). Broad-scale inventories of garden ponds are largely missing in other parts of the world, hampering our understanding of the ecological function of these secondary habitats.

Pond density is a major determinant of aquatic species richness in urban landscapes (Gledhill et al., 2008). Hence, garden ponds have the potential to act as important stepping stones to facilitate the movement of species (e.g., insects, amphibians) across urban landscapes, in addition to increasing freshwater habitat where space is limited (Hill et al., 2021). At the same time, garden and urban ponds in general might be heavily influenced by anthropogenic pressure in the form of multiple types of local management (Blicharska et al., 2016). While most research on garden ponds has focussed on biodiversity (e.g., Hamer and Parris, 2011; Hill and Wood, 2014), there has been no examination of broad-scale patterns in pond design or management practices, or in the spatial distribution of garden ponds at a national level. This is despite the increasing popularity of garden ponds, especially in urban areas where there is limited space. Studies of urban ponds have highlighted the important roles of design and management in shaping aquatic communities (e.g., Blicharska et al., 2016; Oertli and Parris, 2019), yet we have little information on how garden ponds are designed, built and managed. Moreover, there is little information on the age and spatial distribution of garden ponds at multiple spatial scales including whether ponds are located in cities, suburbs or rural districts. This has strong implications for conservation, as garden ponds have high potential to limit future urban biodiversity loss (Hill et al., 2021), although spatial isolation may reduce the ability of ponds to sustain species-rich aquatic communities (Thornhill et al., 2018).

Garden ponds differ from natural ponds in that they generally hold water all year round and are actively managed (e.g., removal of vegetation and silt) that reduces habitat heterogeneity and prevents ecological succession (Biggs et al., 1994; Gaston et al., 2005). They are frequently stocked with fish (Hassall, 2014), and this might determine the design and management types from the onset, including certain decisions on size or cleaning. Therefore, ponds with fish might be distinctive from fishless garden ponds, eventually also leading to different local environments for aquatic species. However, the extent of these potential relationships has not been fully explored.

Small waterbodies are frequently difficult to survey in terms of time and available resources due to their sheer numbers (Kelly-Quinn et al., 2023). The study of garden ponds is further hampered as they often occur on private properties. Therefore, we applied a citizen science approach to inventory garden ponds across Hungary and to collect information from pond owners on pond locations and size, and when ponds were installed and what material was used in their creation. This methodology was particularly crucial given the difficulty for researchers to obtain permission to access garden ponds (Wood et al., 2003). Using an online questionnaire, we also asked pond owners about pond features including management practices and whether ponds were stocked with fish. This way, we could collect a large-scale, country-wide data set on these habitats hidden from the public, but potentially important for aquatic wildlife. While regional inventories of garden ponds have been compiled in western Europe (see Davies et al., 2009), this is the first study to examine garden ponds in central and eastern Europe at a national scale. Our aim was to determine the physical features of a typical garden pond in Hungary, and whether there are any general patterns in these features. We hypothesised that some differences might arise between garden ponds in the heavily populated capital city and ponds in the countryside, possibly originating from differences in preferences of the pond owners. Because the construction of garden ponds has been ongoing for decades, these features and management choices might have also changed with time, for which pond age would be a reliable indicator. In addition to the year and location of construction, key forms of management such as the introduction of ornamental fish might also influence these features and induce changes in other related management practices.

## 2. METHODS

### 2.1. Online survey

In 2021, we commenced the MyPond project (www.mypond.hu) with the broad aims to investigate the biodiversity of garden ponds and if decisions made by pond owners can produce patterns in pond design and management practices across a large geographical extent (i.e., throughout Hungary). In line with these aims, we launched an online survey on the website. The survey consisted of 31 questions related to motivation for building a pond, physical characteristics, management practices (e.g., chemical use, leaf and sediment removal, pond draining), fish and other animals introduced to the pond, and sightings of wild animals (dragonflies, amphibians, birds). The web link was promoted through both traditional and social media outlets. A wide range of media coverage including online and printed articles, TV and radio interviews helped to increase public involvement. We received 834 entries for the survey from 06/07/2021 − 07/09/2022. In assessing patterns and correlations among the variables in this study, we only included pond owners’ responses pertaining to pond location and design (size, construction materials/ substrate type, plantings), fish stocking and management practices (see Table S1 Supplementary Information and Márton et al. in prep.).

### 2.2. Data management

We filtered the data set and excluded ponds that did not match certain criteria. While ponds are often defined as being < 2 ha in area (Hill et al., 2018), we only included garden ponds ≤ 200 m^2^ for further analysis. We based our decision on the observed distribution of pond areas, with the largest pond being 2400 m^2^, while most ponds > 200 m^2^ were agricultural waterbodies. We also excluded ponds if geographical coordinates, length, width or age were not provided. Pond surface area was calculated from lengths and widths provided by pond owners. We took the mean in instances when owners provided a range of lengths or widths. As the ponds can have many different irregular shapes, we opted for a simple multiplication of these two measures. We acknowledge that this might somewhat overestimate sizes but we chose this as a simple and hence reliable type of information our respondents could easily approximate in the form of the widest and longest size of their pond. We used the maximum water depth provided by pond owners. Regarding pond age, if owners did not provide a specific year of installation but stated that they inherited the pond from a previous owner and provided the year they moved in, then pond age was calculated from that year. Conversely, if the current owners said they have lived at the dwelling for example, two years, but entered that the pond was older than 10 years, we retained the pond age as > 10 years. We recorded pond age as ‘0’ if a pond was constructed in the year when the questionnaire was filled out. Pond substrate types were grouped into five categories (concrete, PVC rubber, plastic, metal, natural), with pond descriptions matched as closely as possible to each category. For instance, ‘natural’ substrate included clay and earthen-lined ponds, while ‘plastic’ ponds included fibreglass, polyethylene and polycarbonate materials, and ‘concrete’ included stone and tile ponds. Duplicate data (i.e., pond) records in the questionnaire were omitted.

### 2.3. Landscape variables

According to our original hypothesis that garden ponds might differ based on their location, we used two types of predictors to track these potential differences. First, we plotted the spatial location of each garden pond and determined which were located within the Budapest metropolitan area and those that were outside. The Budapest metropolitan area hosts one-quarter of the population and is the most developed region of Hungary (Egedy et al., 2017). This binary predictor was included in the analyses as “Budapest”. Second, we calculated the area of urban land and wetlands within a 1-km radius around each garden pond using the Ecosystem Map of Hungary to classify land cover types (project KEHOP-430-VEKOP-15-2016-00001, Ministry of Agriculture, 2019). A 1-km radius was chosen as it encompasses the landscape influences in many urban pond communities (Oertli and Parris, 2019). Urban land contained pixels of tall buildings, short buildings, sealed roads, railways and other artificial surfaces, whereas the wetlands category contained pixels of wetlands, marshlands, temporary wetlands, lakes and other standing waters. The number of raster pixels (20 x 20 m) in the Ecosystem Map covered by the urban and wetlands categories were summed within a 1-km radius circle around each pond, and the proportion cover of each was calculated. All spatial analyses were conducted using QGIS 3.28.1 (QGIS Development Team, 2022).

### 2.4. Statistical analysis

Given that our set of 21 variables included a mix of continuous (latitude, longitude, area, depth, urban land cover and wetland cover), ordinal (age) and binary variables (Budapest vs. countryside, four possible substrate types, introduction of fish, presence of shoreline vegetation, presence of aquatic vegetation, and application of six types of possible pond management practices), we opted for a series of complementary analyses to track relationships among them.

#### 2.4.1. General patterns and correlations

First, we assessed correlations among the variables (excluding site latitude and longitude) using Spearman’s rank correlations (*r_s_*). Then, a Multiple Factor Analysis (MFA) was conducted which can combine continuous variables with binary data in a non-constrained ordination, and in case of categorical variables, it can also consider the group structure and combine information on each category. MFA is an extension of Principal Component Analysis (PCA) used to analyse several data sets measured on the same objects, and provides a set of common factor scores (Abdi et al., 2013). The MFA was used to visualise general patterns and associations among 19 variables structured into eight groups: (1) pond location, i.e., within/outside Budapest, (2) urban/wetland cover, (3) pond area, (4) pond age, (5) pond substrate type, (6) presence of aquatic and shoreline vegetation, (7) fish introduction, (8) management practices. Groups 2 and 3 consisted of quantitative variables; while groups 1 and 5 – 8 were qualitative variables. We excluded pond depth from the analysis and kept a single measure of pond size, as pond area and depth were strongly correlated (*r_s_* = 0.483; Table S2). Pond age was entered on an ordinal scale from 0 – 10 years and > 10 years, with a value of ‘0’ indicating that the pond had been created in the year of filling out the survey questionnaire. All other qualitative variables were entered into the analysis as binary variables, with a ‘1’ confirming presence at a pond. Continuous variables were standardised by scaling to unit variance. We conducted the MFA in program R v4.3.0 using the packages FactoMineR (Husson et al., 2023) and factoextra (Kassambara and Mundt, 2020).

Finally, we converted all non-binary variables to binary data (by binning continuous variables and creating dummies from the categorical variables), and analysed co-occurrence patterns among the variables (excluding latitude and longitude). We transformed the continuous variables into categorical variables based on size distributions, each with three classes (area: small < 2 m^2^, mid 2 – 20 m^2^, large > 20 m^2^; urban: low < 0.3, mid 0.3 – 0.6, high > 0.6; age: new < 2 yrs, mid 2 – 10 yrs, old > 10 yrs). We used a probabilistic model to test for statistically significant patterns among pond variables. The model gives the probability that two variables would co-occur at a frequency less than (or greater than) the observed frequency if the two variables were distributed independently of one another, and classifies associations as negative, positive or random (Veech, 2013). We used the cooccur R package (Griffith et al., 2016) on a final dataset of 24 variables.

#### 2.4.2. Role of spatial position, pond age, and the introduction of fish

To test for the possible roles of spatial location (geographical coordinates, urban land cover), age, and whether the pond was used for keeping ornamental fish, we applied a further three methods: Mantel correlograms, distance-based Redundancy Analysis and Moran’s Eigenvector Maps. We first tested for the spatial autocorrelation in the similarity of pond features by means of a Mantel correlogram (Oden and Sokal, 1986), using the ordination scores from the MFA, to be able to test for potentially similar trends in pond design in local geographical areas. We used the vegan (Oksanen et al., 2022) and fields (Nychka et al., 2022) R packages to relate the component scores derived from the MFA for each pond on dimensions 1 and 2, to pond geographical position (latitude, longitude). We initially calculated pairwise geographic Euclidean distances between all ponds. The computed pairwise distances between component scores and log-transformed spatial distances were used to perform a Mantel test with 999 permutations to calculate Mantel correlation coefficients for ten distance classes.

We then also directly tested for the relative effect of age, presence of fish, urban land cover, and eigenvector-based spatial structure on the 19 measured variables using distance-based Redundancy Analysis (dbRDA) and subsequent variation partitioning. In the dbRDA, we used predictor variables that either contributed the greatest variation on dim1 and dim2 in the MFA (presence of fish, urban land cover), or were plausible drivers of why pond owners would select specific pond designs and implement certain management practices (age), thereby resulting in patterns in the explanatory variables. We also included the eigenvectors from Moran’s Eigenvector Maps (MEM; Dray et al. 2006) that best modelled positive spatial correlation of the longitude and latitude of the garden pond locations. We included the MEMs as spatial explanatory variables in the dbRDA to identify potential spatial structuring within the pond locations (see below). Among the pond features, pond area was log_10_ (-x×10) transformed while the square root of the proportion of urban land cover was taken to normalise the data before including them in the dbRDA model. We also converted pond age into two dummy variables (corresponding to the two most contrasting categories): (1) newly built ponds < 6 years old; and (2) older ponds > 10 years since installation. The other categorical variable (substrate type) was also converted to a binary (dummy) variable; we used the four most frequent pond substrate types and omitted the one with only three occurrences (metal) to keep the most representative types on the ordination plot.

We conducted the dbRDA with Euclidean distance using 999 permutations in vegan using the capscale function; i.e., the dissimilarity data are first ordinated using metric scaling, and the ordination results are analysed as an RDA (Legendre and Anderson, 1999). In order to identify the best model and individual predictor variables that significantly explained the variance in the explanatory variables we used the ordistep procedure in vegan, using both forward and backward stepwise model selection and a maximum of 200 permutation tests.

The ordistep function performs step-wise selection of environmental variables based on two criteria: if their inclusion into the model leads to a significant increase of explained variance and if the AIC of the new model is lower than AIC of the simpler model.

We constructed MEMs using the adespatial (Dray et al., 2023) and adegraphics R packages (Dray and Siberchicot, 2023). The MEM method consists of identifying a binary connectivity matrix defining which pairs of sites are connected and which are not, and a weighting matrix providing the intensity of the connections (Borcard et al., 2011). We computed eigenvectors of a spatially weighted matrix of garden pond latitude and longitude (Borcard and Legendre, 2002). Using the ordistep procedure based on F and p values, we determined which of the first 20 MEM eigenvectors (MEM1 – MEM20) best explained the variation in the pond variables, using both forward and backward stepwise model selection and a permutation maximum of 999. We did not use all MEM eigenvectors in the null model as we were only interested in large-scale regional differences, which are best modelled with those MEM eigenvectors with the highest eigenfunctions (and hence highest rank). The two MEMs with the highest explanatory power and urban land cover, fish presence and pond age (young or old) were included as predictor variables in the final dbRDA model.

Subsequently, we used variation partitioning in the dbRDAs to determine the relative importance of the spatial (two best MEMs) and three environmental variables (urban land cover, age, fish) versus 13 explanatory variables grouped under pond area, substrate, vegetation and management. Variation partitioning aims to quantify the various unique and combined fractions of variation explained by two subsets of variables (Borcard and Legendre, 2002). We used a Monte Carlo permutation test for capscale under a reduced model (999 permutations) performed with the anova function, with the subsets assessed on F statistics.

## 3. RESULTS

### 3.1. General pond features

A total of 753 garden ponds were retained for further analysis, spread across Hungary (Fig. 1), with 127 ponds (17%) located within the Budapest metropolitan area and 626 (83%) located outside the capital city. The mean pond surface area was 18.7 m^2^ and biased towards small pond size classes (Table 1; Fig. 2a). The total surface area of the 753 garden ponds was 14,064 m^2^ (1.4 ha). The mean proportions of urban land and wetlands within a 1-km radius around a pond were 0.284 and 0.023, respectively, with substantial range in values (Table 1). The mean distance to the nearest garden pond across the country was 3514 m, although garden ponds located in Budapest were substantially closer to one another (mean = 735 m), while ponds outside the capital were substantially further apart (mean = 4089 m; Table 1).

**Fig. 1.**
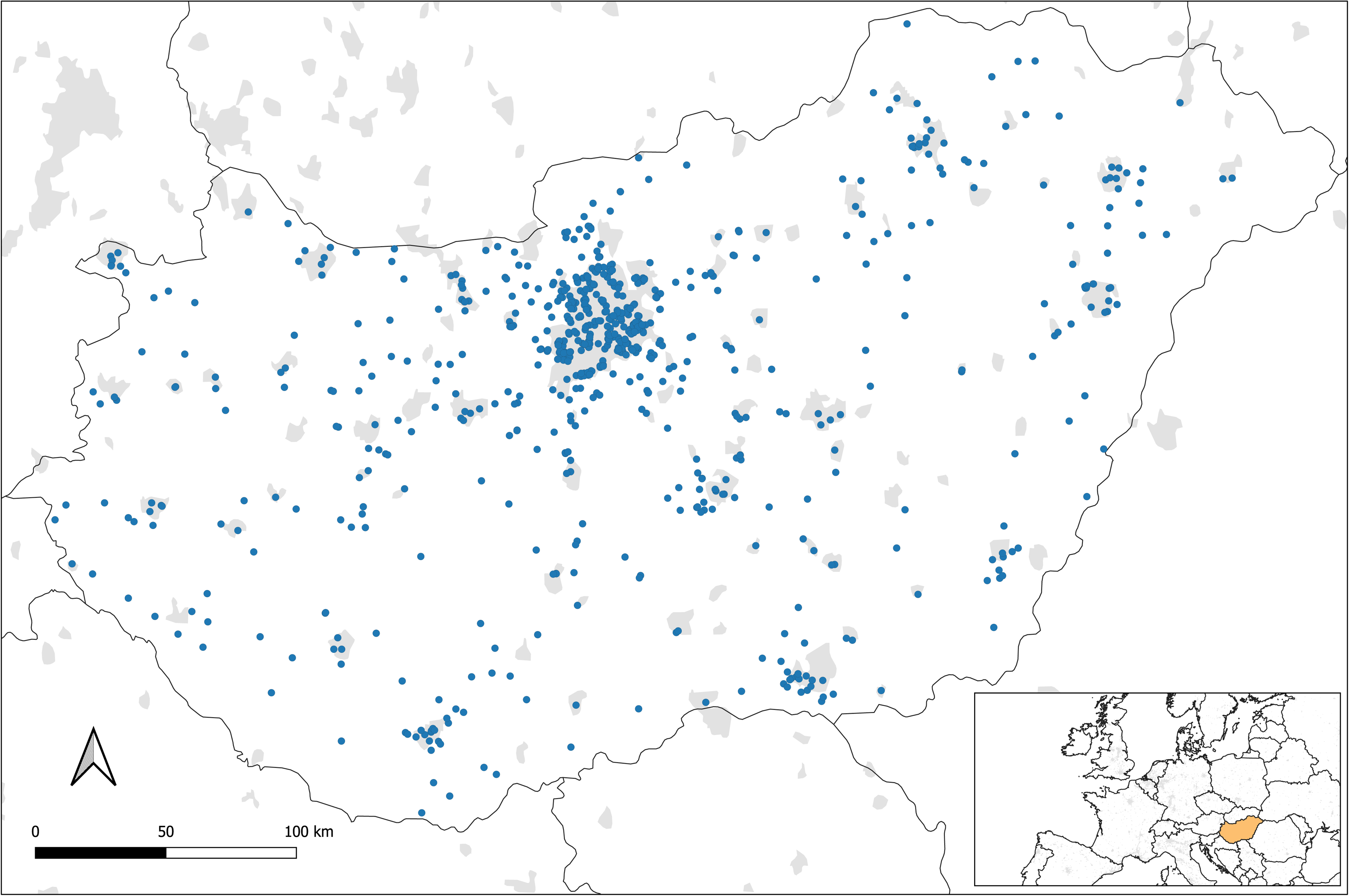
Map of the distribution of 753 garden ponds for which responses were received in the MyPond survey questionnaire, Hungary. Urban areas are shown in grey (derived from 2002-2003 MODIS satellite data at 1 km resolution; Schneider et al., 2003). The capital city (Budapest) is at the top centre of the map (high pond density).

**Fig. 2.**
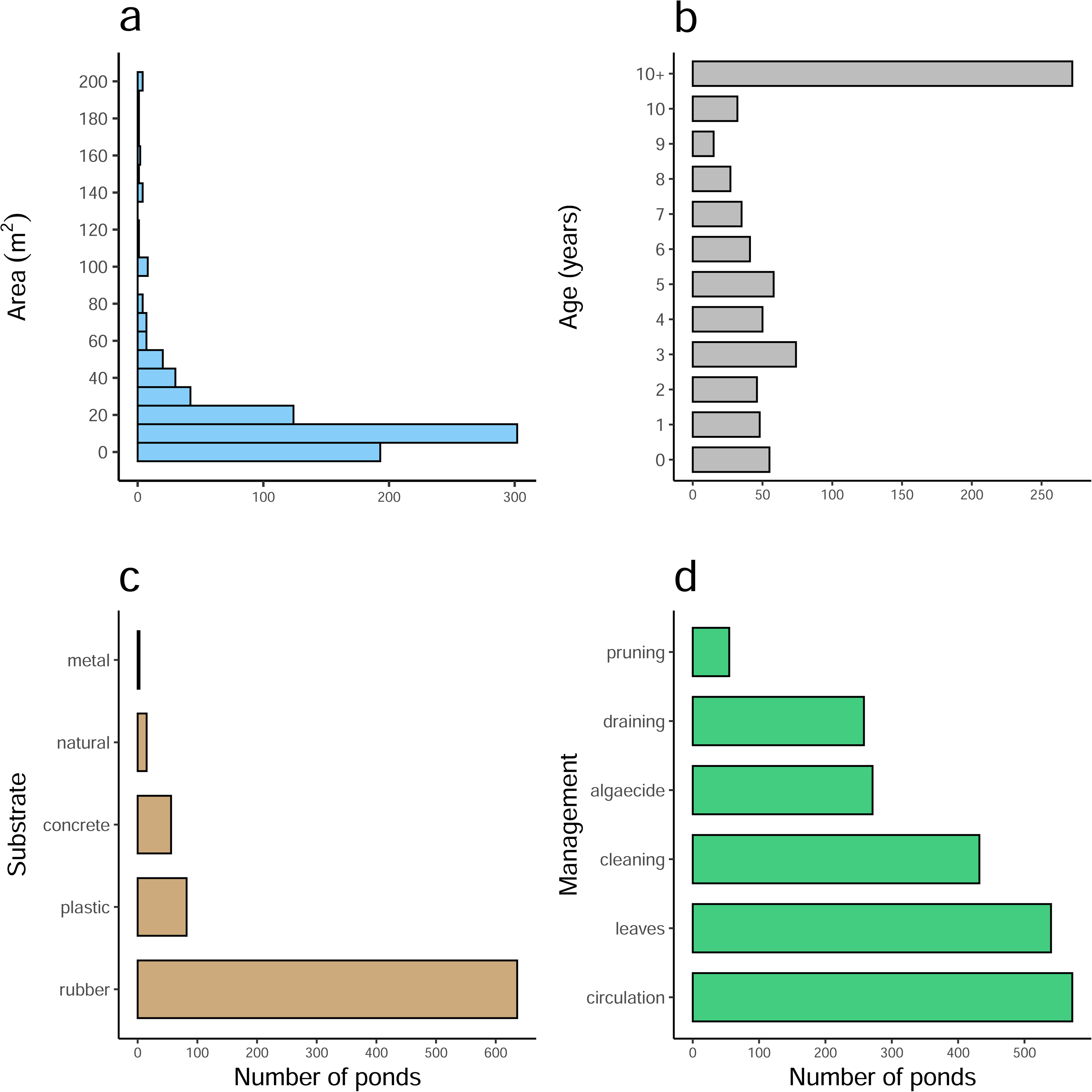
Number of 753 garden ponds according to (a) area, (b) age (years since installation or construction), (c) substrate type, and (d) management practice (see Table S2 for an explanation of variable names).

**Table 1.** Descriptive statistics for continuous variables recorded at 753 garden ponds in Hungary.

| **Variable** | **mean** | **SD** | **min** | **max** |
| --- | --- | --- | --- | --- |
| Area (m<sup>2</sup>) | 18.68 | 27.36 | 0.25 | 200.00 |
| Depth (cm) | 120.65 | 45.96 | 10.00 | 450.00 |
| Urban | 0.284 | 0.210 | 0.000 | 0.974 |
| Wetlands | 0.023 | 0.046 | 0.000 | 0.421 |
| Distance (m) to nearest pond (Hungary) | 3514 | 4518 | 4 | 29063 |
| Distance (m) to nearest pond (Budapest) | 735 | 566 | 68 | 3229 |
| Distance (m) to nearest pond (ex-Budapest) | 4089 | 4751 | 4 | 29063 |

Area = pond surface area; depth = maximum water depth; urban = proportion of urban land cover within a 1-km radius of a pond; wetlands = proportion of wetlands and waterbodies within a 1-km radius of a pond; Distance to nearest pond (Hungary) = distance to the nearest pond considering all ponds in Hungary; Distance to nearest pond (Budapest) = distance to the nearest pond considering only ponds located within the Budapest metropolitan area; Distance to nearest pond (ex-Budapest) = distance to the nearest pond considering only ponds located outside of the Budapest metropolitan area

Thirty-six percent of ponds were constructed over a decade ago while 7% were new ponds (<1 year old) and 64% were ≤10 years old (Fig. 2b). The majority of ponds (84%) were constructed of rubber (PVC) lining (Fig. 2c). Only three ponds were built with metal (e.g., tin) while 15 ponds had natural earth substrate (Fig. 2c). Thirty-eight ponds (5%) were constructed of more than one substrate type; e.g., 28 ponds had a mix of concrete and rubber lining. Most ponds were planted with aquatic and shoreline vegetation (93% and 84%, respectively) and fish had been introduced in 85% of ponds. In terms of management practices, 76% of ponds had a water circulation device installed, while leaves were actively removed by pond owners from 72% of ponds (Fig. 2d). Thirty-four percent of ponds were drained to remove sediment that had accumulated on the pond bed (Fig. 2d). Chemical algaecides were used more frequently than natural probiotic algaecides (33% and 3% of ponds, respectively).

### 3.2. Correlations and associations between pond location, design and management

There were no strong correlations between the proportion of urban land cover within a 1-km radius and either pond surface area (*r_s_* = –0.172) or the proportion of wetlands (*r_s_* = –0.364; Table S2). There were moderately strong correlations between pond area and water depth (*r_s_* = 0.483; Table S2), and between ponds in Budapest and the proportion of urban land cover (*r_s_* = 0.496; Table S2). There was a positive correlation between pond cleaning (sediment removal) and draining (*r_s_* = 0.396; Table S2). There were no strong correlations among the other variables (*|r_s_|* < 0.5; Table S2).

The first dimension of the MFA (dim1) explained 7.43% of the variance in the data of the 753 ponds while the second dimension (dim2) explained 6.98% (Fig. 3a). Dominant contributions to dim1 and dim2 were fish introductions at a pond (28.3%) and ponds located in Budapest (32.4%), respectively (Figs. 3b and 3c). Pond vegetation and management also contributed substantially to dim1 (18 – 20%) while pond landscape contributed substantially to dim2 (31.7%). There was no substantial contribution of pond age to either dimension (Fig. 3a).

**Fig. 3.**
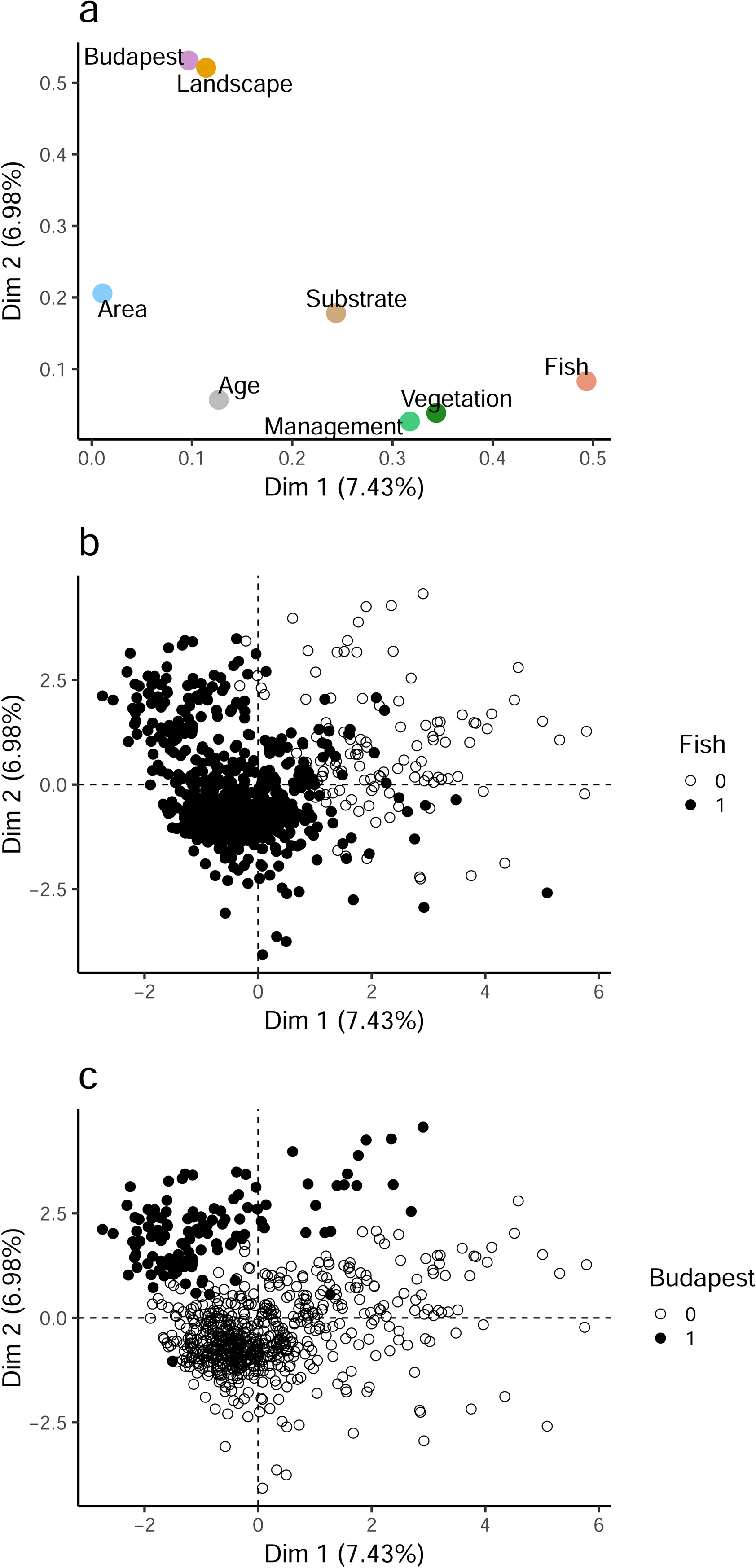
(a) Two-dimensional plot of the multiple factor analysis (MFA) performed on 19 variables (8 groups) recorded from 753 garden ponds in Hungary; pond coordinates on MFA dimensions grouped by: (b) introduction of fish, and (c) ponds located in Budapest.

Co-occurrence patterns revealed 59 significantly positive and 60 significantly negative associations among the 24 variables. The stocking of fish at a pond had nine and five significantly positive and negative associations with other variables, respectively (Fig. 4). Fish were more likely to have been introduced in ponds > 2 m^2^, aged 2 – 10 years, lined with rubber, with aquatic and shoreline vegetation, and being actively managed (e.g., via sediment and leaf removal, algaecide application, draining) with water circulation devices installed (Fig. 4). Multiple management practices were positively associated with each other (Fig. 4). The oldest ponds tended to be the largest and were mostly located at intermediate levels of urbanisation. The newest ponds were associated with low levels of urbanisation, they were less likely to be managed by draining or cleaning, and while they were less likely to contain fish, the application of algaecides was more likely. High urbanisation was associated with a higher likelihood of a plastic pond (Fig. 4).

**Fig. 4.**
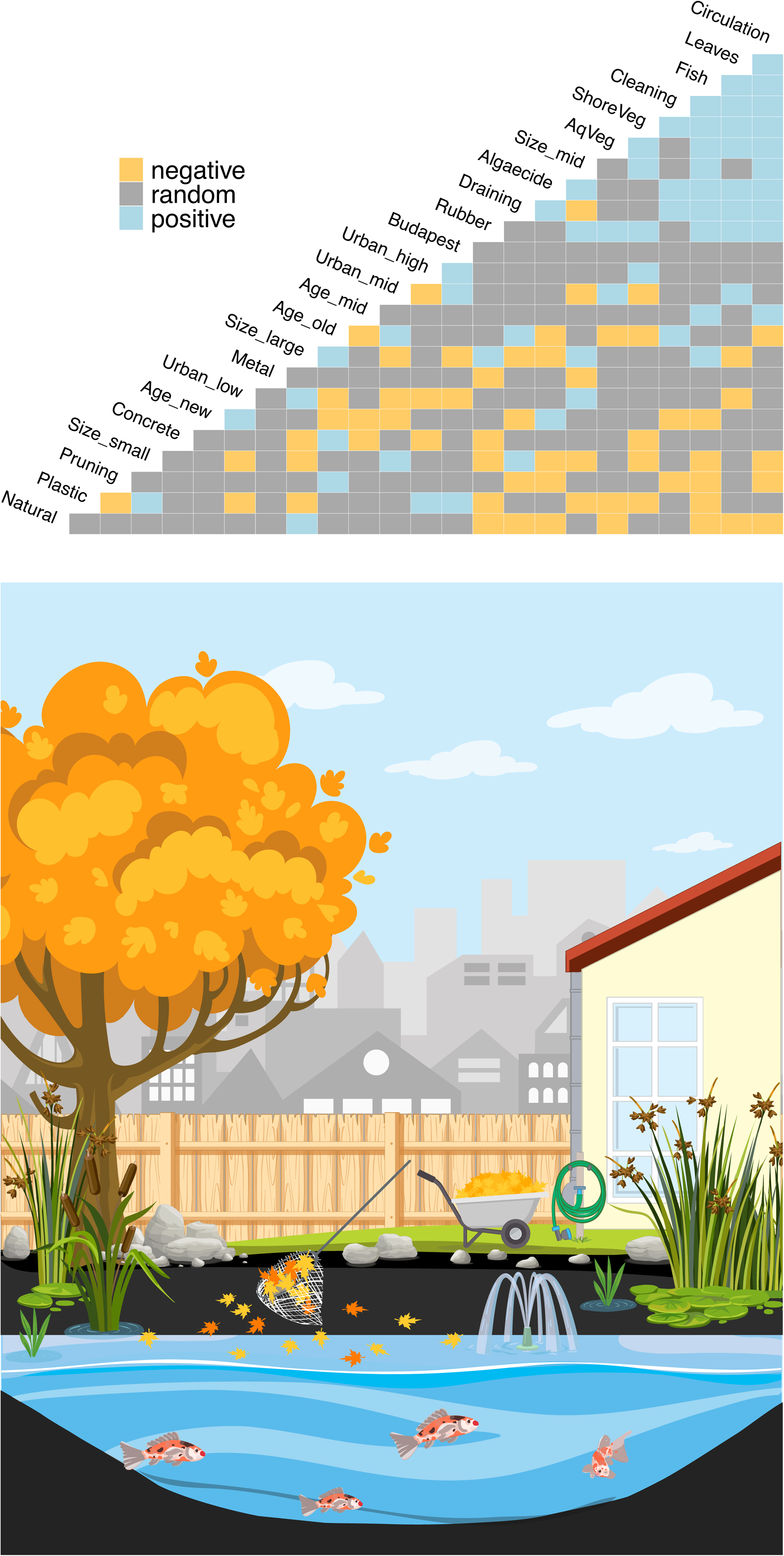
Heat map showing the significantly positive and negative variable associations determined by the probabilistic co-occurrence model for 753 garden ponds in Hungary. (See Table S2 for a description of the variables). Pond size, age and urban land cover within a 1-km radius of a pond were each converted to three categories. A typical garden pond based on the most frequent positive associations is depicted (see Fig. S2 for a photograph of a typical garden pond).

### 3.3. Geographical differences in pond design and management

There was a significant distance-decay relationship for dim1 and dim2 of the MFA (Fig. 5). There was positive spatial autocorrelation with both dimensions among garden ponds at distances up to 73 km, with significant relationships at 15 km (Mantel test: *r* = 0.670, p = 0.001) and 44 km (*r* = 0.293, p = 0.002; Fig. 5). The distance-decay relationship showed significantly negative autocorrelation among ponds in all distance categories above 73 km (Fig. 5).

**Fig. 5.**
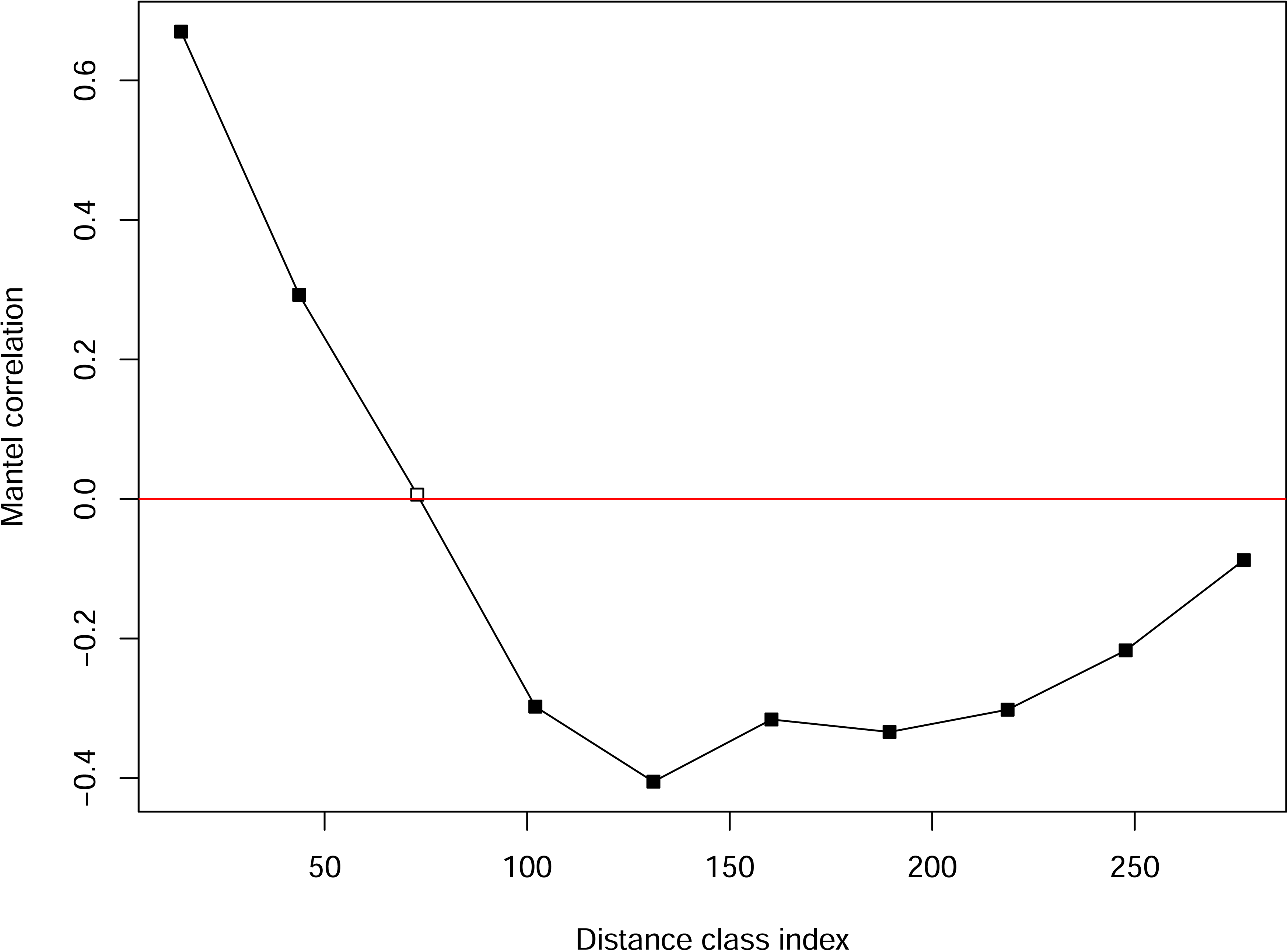
Mantel correlogram for MFA dimensions 1 and 2 and geographic distance (log-transformed) between 753 garden ponds. Closed boxes indicate significant correlations within a distance class, tested within ten distance classes. Values above the red line indicate positive spatial autocorrelation in dim1 and dim2 among ponds.

Two eigenvector maps were included in the best model: MEM1 (AIC = 822.6; F = 8.51) and MEM3 (AIC = 816.8; F = 2.65), which represented a separation of ponds between the capital city and countryside (MEM1), but also within Budapest itself (MEM3) with ponds divided between the southern and northern city districts (Fig. S1).

### 3.4. Spatially-structured relationships between pond location, design and management

The first two eigenvalues of the dbRDA explained 93% of the total variation in the explanatory variables, with the first axis (CAP1) contributing the greatest in terms of the constrained eigenvalues (CAP1: 0.186; CAP2: 0.026). Like the MFA, the first axis of the biplot described a gradient of garden ponds with or without fish present (Fig. 6). The second axis described a gradient of ponds surrounded by high urban land cover to old ponds distributed outside the capital city (Fig. 6). The permutation test of the dbRDA showed that fish presence explained most of the variance in the model (F = 38.13, variance = 0.141; Table 2) while urban land cover also explained significant variation (F = 15.23, variance = 0.056; Table 2). The two MEMs contributed little variation to the model (MEM1: 0.007; MEM3: 0.008; Table 2).

**Fig. 6.**
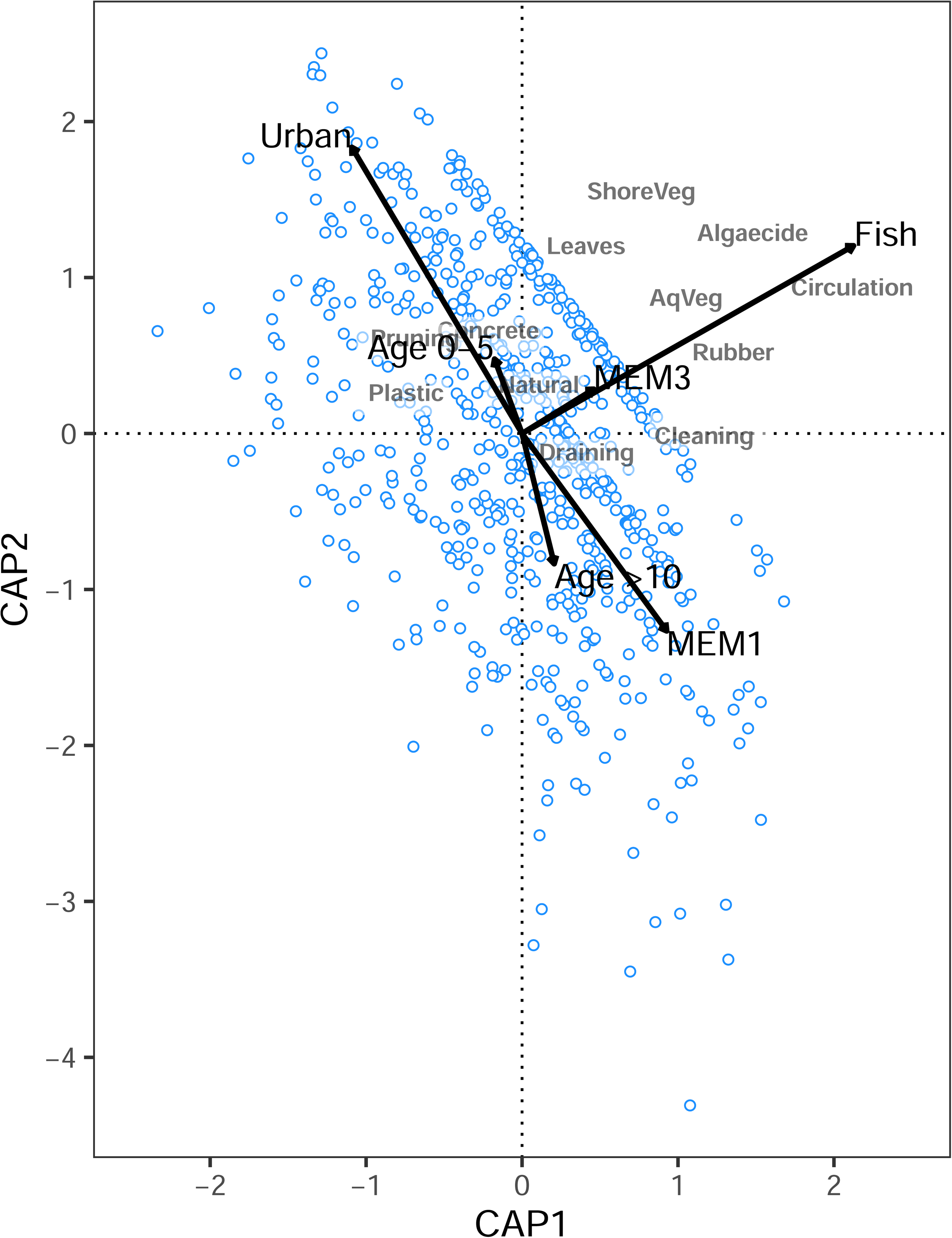
Distance-based redundancy analysis (dbRDA) biplot of six predictor variables and 13 explanatory variables. See Tables 2 and S2 for a description of predictor and explanatory variables, respectively.

**Table 2.** Results of the Monte Carlo permutation test for the distance-based redundancy data analysis (dbRDA) for 753 garden ponds in Hungary. d.f. = degrees of freedom; var = variance; *significant (p < 0.05).

| Predictor | d.f. | var | F | Pr(>F) |
| --- | --- | --- | --- | --- |
| Fish | 1 | 0.141 | 38.13 | 0.001* |
| Urban | 1 | 0.056 | 15.23 | 0.001* |
| Age (<6 yrs) | 1 | 0.005 | 1.37 | 0.199 |
| Age (>10 yrs) | 1 | 0.011 | 3.04 | 0.025* |
| MEM1 | 1 | 0.007 | 1.87 | 0.102 |
| MEM3 | 1 | 0.008 | 2.17 | 0.066 |
| Residual | 746 | 2.752 | - | - |

The predictor variables (urban land, age, fish) could explain more variation in the explanatory variables than the purely spatial variables (MEMs), although overall only a very small proportion of the unique variance was explained (environmental predictors: 0.06; MEMs: 0.00). The RDA model of the purely environmental predictors (F = 12.50; p = 0.001) and the model of the spatial variables (F = 2.02; p = 0.05) were both significant.

## 4. DISCUSSION

Garden ponds may provide small patches of supplementary freshwater habitat in areas where urbanisation and agricultural intensity have caused widespread pond loss (Gibbons et al., 2023; Gledhill et al., 2008). Hence, they may increase the availability of ponds for aquatic taxa and contribute to landscape connectivity as stepping-stones (Hill et al., 2021). Our country-wide citizen science survey, based on responses from 753 pond owners, revealed considerable variation in pond size (0.25 – 200 m^2^), and that most ponds were <10 years old, were rubber-lined with water circulation (e.g., fountains), and had fish introduced together with aquatic and shoreline vegetation. The introduction of fish was the strongest driver of the other local factors and accounted for the greatest amount of variation, followed by pond geographic location (i.e., the capital city, Budapest, or elsewhere).

### 4.1. Fish, pond size and design

Survey respondents reported they had introduced fish in 85% of the garden ponds. Subsequently, the deliberate stocking of fish was a significant driver of variables related to pond design and management. Keeping ornamental fish is a popular and widespread practice in central Europe (Patoka et al., 2017), and likely to the primary motivation to build a pond (Hill et al., 2021). Fish introductions were also associated with older, larger and deeper ponds. Fish species identity will likely influence the decision made on pond size – for example, koi carp will require a larger area than the smaller ornamental varieties (e.g., goldfish). New ponds (<2 years old) were less likely to have fish introduced. This may be because pond owners wanted to stabilise them before introducing fish, or may reflect a recent trend to have fishless garden ponds to support native biodiversity.

The average size of the ponds in our study (18.7 m^2^) is considerably larger than that reported for garden ponds in the United Kingdom, where similar data are available. There, smaller mean sizes were found at multiple spatial scales (national survey: 1 m^2^ in Davies et al., 2009; regional surveys: 2.53 m^2^ in Gaston et al., 2005; 5.0 m^2^ in Hill et al., 2021). This concurs with a general difference in the ratio of garden ponds stocked with fish in Hungary (85%; this survey) and in the UK (approx. 30%; Loram et al., 2011).

Ultimately, pond size is likely to be determined by the availability of space in a resident’s garden, and highly urbanised areas often only contain small ponds due to limited space (Oertli et al., 2023). In the UK, garden ponds generally are more likely to be found in large than in small gardens (Loram et al., 2011). In our case, residents in Budapest may have limited space available and so install smaller plastic ponds, as the mean pond area for Budapest was 12.5 m^2^ while elsewhere it was 19.9 m^2^.

Pond size is an important driver of biodiversity in urban landscapes as larger ponds tend to host higher species richness (Oertli and Parris 2019). However, in our dataset, larger ponds tended to be the ones with fish, which can decrease the richness and abundance of multiple pond taxa (Trovillion et al. 2023). The relatively small size of garden ponds compared to other urban ponds may limit their ability to support highly diverse aquatic communities at the local scale, and may lack habitats important for specific taxa such as dragonflies (e.g., emergent vegetation; Hill et al., 2021). At the same time, 93% of our surveyed ponds contained aquatic vegetation, suggesting a relatively large amount of heterogeneity in habitat structure, and therefore could support biodiversity across broad spatial scales. Vegetation in urban ponds provides shelter and refuge for a range of organisms (e.g., turtles, amphibians and invertebrates; Hamer and McDonnell, 2008; Marchand and Litvaitis, 2004; Thornhill et al., 2017). This, together with the relatively larger mean size may increase the chances of garden ponds in Hungary to provide viable freshwater habitats for a variety of taxa. For example, respondents of the same survey regularly saw birds, amphibians and dragonflies in their ponds (Márton et al., in prep.).

We found that most garden ponds were lined with PVC rubber or made of plastic. Owners likely select such designs to ensure consistently high water levels, especially in ponds with ornamental fish. While there is considerable flexibility in the possible size, shape and depth of rubber-lined ponds, prefabricated plastic ponds have pre-determined dimensions, and tend to be overall smaller. For example, a major gardening retailer in Hungary sells prefabricated polyethylene ponds as “quick and convenient garden ponds”, with dimensions 135 × 105 × 45 cm (length × width × depth), although larger models were also available (265 × 225 × 80 cm). These small plastic ponds are more typical for ponds in the Budapest metropolitan area, under high levels of urbanisation. While they were more likely to be fishless (likely due to their small size), they also lacked vegetation according to our results, which makes their role for biodiversity difficult to assess. Ornamental ponds with an artificial bed such as PVC liner or concrete tend to have lower biodiversity than urban ponds with a natural substrate (Oertli et al., 2023).

### 4.2. Pond management

There was a wide range of management practices applied at the garden ponds, with multiple practices often employed. Most ponds had a water circulation device installed to enhance water quality through increased dissolved oxygen and water transparency (Hao et al., 2021), but also for aesthetic purposes (e.g., ornamental fountains). These features can have positive outcomes for pond biodiversity, as they provide a relatively unpolluted aquatic habitat within urban areas (McGoff et al., 2017), and the increasing popularity of garden ponds in Hungary is likely to contribute to relatively clean sources of freshwater for wildlife in residential areas. Other common management practices such as the removal of leaves, however, may decrease habitat heterogeneity for benthic organisms (Biggs et al., 1994). At the same time, the retention of bottom sediment and planted vegetation might compensate for any reduction in habitat structure (Soukup et al., 2022). Hence, pond cleaning may have different outcomes depending on the taxa. Pond owners also tended to keep their garden ponds neat for aesthetic purposes, by cutting and removing vegetation, which can subsequently reduce habitat diversity and species richness (Noble and Hassall, 2015; Oertli and Parris, 2019). Chemical algaecides, rather than “natural” probiotic algaecides, were used more frequently in ponds.

An analysis of the occurrence of animal taxa in our data set found a negative relationship between amphibians and their larvae and algaecide usage (Márton et al. in prep.). Chemical algaecides decrease tadpole survival and growth rates most likely due to copper contaminants (Christenson et al., 2014). In line with this, an analysis of the occurrence of animal taxa in our data set found a negative relationship between amphibians and their larvae and algaecide usage (Márton et al. in prep.). Overall, the wide range and frequency of management practices in garden ponds may be a major driver of aquatic communities in urban areas (Hill et al., 2021).

Pond management may change with new fashions or different owners, and together with pond turnover, has the potential to affect local biodiversity (Gaston et al., 2005). For example, new species of ornamental plants or novel cleaning methods including types of chemicals used may become popular. New trends in management may also be encouraged by high profile celebrity gardeners (Hassall et al., 2016). In our data, we did not observe strong temporal trends in overall management practices; however, we found that the newest ponds were less managed and were in less urbanised landscapes. Newer ponds were also less likely to be cleaned out, which could either indicate a potential change in the perception of pond management, or that newer ponds had not yet accumulated large amounts of sediment.

### 4.3. Pond landscapes

There was evidence of spatial structuring according to the geographical location of garden ponds, with a separation between ponds in Budapest and those located outside the capital. This pattern could imply there is a greater density of ponds within the city compared to other parts of Hungary. However, city dwellers may have had greater interest in the questionnaire, potentially leading to a bias towards city ponds and therefore higher sampling intensity in Budapest. Nonetheless, a higher pond density in Budapest would correspond to high pond densities (Gaston et al., 2005) and positive spatial autocorrelation observed in other European cities at roughly similar spatial scales to what we found (Hill et al., 2017). Despite the spatial structure evident, we found greater importance of fish and urbanisation in explaining variation in the data than purely spatial effects, highlighting the importance of landscape context and pond owner motivations (e.g., to have ornamental fish) in how garden ponds are designed and managed.

Most garden ponds were surrounded by high levels of urban land cover (e.g., buildings, roads), which can reduce habitat connectivity and have a negative effect on species richness of macroinvertebrate and amphibian communities (Blicharska et al., 2017; Hamer and Parris, 2011; Heino et al., 2017). Most ponds also had a very low percentage (2.3%) of wetlands within a 1-km radius, indicating that garden ponds may be the only aquatic habitat available in many areas. This finding has important implications in maintaining landscape connectivity for pond-dependent fauna, as garden ponds may act as stepping-stones for movement and as refuges for aquatic taxa in otherwise largely inhospitable landscape settings (Gledhill et al., 2008). For instance, the average distance to the nearest garden pond outside of Budapest was 4089 m, which is close to the maximum dispersal distance of many pond-dependent organisms in central and eastern Europe. However, the average inter-pond distance in Budapest was 735 m which is well within the dispersal capabilities of some organisms (Kappes and Haase, 2012; Conrad et al., 1999; Trochet et al., 2014).

### 4.4. Unexplained variation

Despite the clear patterns we observed, the variables that we derived from our questionnaire to pond owners (age, location, fish stocking) had very little explanatory power, indicating that some important drivers of variation were not included in the dataset. Socio-economic factors may be important drivers behind the decisions made by pond owners, although the socio-economic status of residents does not necessarily lead to patterns in terms of the management and function of urban ponds, or subsequent effects on biodiversity (Blicharska et al., 2017). However, more garden ponds are often built in affluent suburban areas where there is more garden space available (Gledhill and James, 2012), yet may be more intensely managed and support less biodiversity (Blicharska et al., 2017).

## 5. CONCLUSION

Using responses from 753 pond owners, we evaluated patterns in garden pond design and management practices, and in pond locations throughout Hungary. We found that pond design was strongly related to the introduction of fish. This pattern indicates that homeowners in Hungary are creating ponds in their backyards primarily to stock ornamental fish, and are landscaping ponds by installing features to maintain water circulation (e.g., fountains) and planting aquatic vegetation while managing ponds by occasionally draining the water and removing sediment and fallen leaves, and applying algaecides. Using spatial statistics, we found differences between garden ponds in Budapest and elsewhere in the country, suggesting there may be differing trends in pond designs, possibly related to the limited area available in city backyards. Furthermore, it is likely that the pond designs we observed reflect relatively recent trends, as we found that most garden ponds had been created in the past decade.

While we have no estimates on the number of garden ponds currently in Hungary, we consider our sample of 753 ponds to be a representative cross-section of the types of garden ponds in Hungary, especially given the broad geographical extent of pond distribution. Even simply obtaining geographical information on the location of garden ponds is essential in monitoring this resource due to their small size and lack of detailed mapping data (Hassall, 2014). We recommend the further documentation of management practices employed at garden ponds to gain insight into how pond owners can maximise the ecological role of these ponds in urban landscapes and benefit biodiversity.

## Declaration of Competing Interest

The authors declare that they have no known competing financial interests or personal relationships that could have appeared to influence the work reported in this paper.

## ACKNOWLEDGEMENTS

This work would not have been possible without the involvement of 834 citizen scientists filling out our survey. We thank Vivien Kardos for help in preparing the garden pond illustration and Kata Bene for assistance in data curation. Virág Csiszár assisted in designing the online survey form. We acknowledge funding from the Eötvös Loránd Research Network (currently Hungarian Research Network), the RRF-2.3.1-21-2022-369-00014 project and the Sustainable Development and Technologies National Programme of the Hungarian Academy of Sciences (FFT NP FTA). Andrew Hamer and Barbara Barta acknowledge further support from OTKA K142296. Barbara Barta was supported by the ÚNKP-22-3 New National Excellence Program of the Ministry for Culture and Innovation from the source of the National Research, Development and Innovation Fund (ÚNKP-22-3-I-ELTE-568). Irene Tornero was supported by the EU NextGenerationEU, Ministry of Universities and Recovery, Transformation and Resilience Plan via Universitat de Girona (REQ2021_A_34). Zsófia Horváth acknowledges further support from the Bolyai+ Grant (ÚNKP-22-5-ELTE-84).

## Notes

### Competing Interest Statement

The authors have declared no competing interest.

