## Supplementary Material for "Patterns and correlates in the distribution, design and management of garden ponds along an urban-rural gradient"

**Table S1.** Questionnaire from the MyPond survey.

Note that only questions relating to topics explored in this paper are included (see Márton et al. in prep. for other questions from the survey).

| **Questions** | **Answer options** |
|---|---|
| What are the coordinates of your garden pond?* | |
| How old is your garden pond? | Made this year ☐<br>1 year old ☐<br>2 years old ☐<br>3 years old ☐<br>4 years old ☐<br>5 years old ☐<br>6 years old ☐<br>7 years old ☐<br>8 years old ☐<br>9 years old ☐<br>10 years old ☐<br>Older than 10 years ☐<br>I do not remember ☐<br>Made by the previous owner, I bought it with the house ☐ |
| If your answer to the previous question was that you had bought the house with an existing pond, please write approx. how many years ago you bought the house! | |
| How deep is the pond (cm)? | |
| What is the pond's length (cm)? | |

| | | |
|---|---|---|
| What is the pond's width (cm)? | | |
| Please check the appropriate answer regarding the substrate of the pond! (Check all that apply! If you answered “Other”, please detail what material you used) | Foil (PVC/rubber)<br>Concrete<br>Plastic basin<br>Other: | □<br>□<br>□<br>□ |
| What kind of introduced animals are there in the pond? (Check all that apply!) | Fish<br>Turtle<br>None<br>Other: | □<br>□<br>□<br>□ |
| Did you plant any plants in the garden pond? | Yes, on the shoreline (sedge, iris, etc.)<br>Yes, in the water (water lily, pondweed, etc.)<br>In both places<br>No | □<br>□<br>□<br>□ |
| Do you drain the water of the pond? | Yes<br>No | □<br>□ |
| Do you clean the pond bed, e.g. to remove the sediment? | Yes<br>No | □<br>□ |
| Do you use any other treatments? (Check all that apply! If you answered "other", please detail what treatment you use!) | Yes, algaecide<br>Yes, I remove the leaves<br>Yes, other<br>No | □<br>□<br>□<br>□ |
| Do you use any equipment to circulate the water (e.g. filter, fountain)? | Yes<br>No | □<br>□ |

*respondents were given instructions on how to extract the coordinates of their garden ponds using Google Maps©.

**Table S2.** Spearman's rank ($r_s$) correlation coefficients among 20 variables recorded at 753 garden ponds, Hungary.

| | Area | Depth | Urban | Wetlands | Budapest | Age | Fish | AqVeg | ShoreVeg |
|---|---|---|---|---|---|---|---|---|---|
| Depth | **0.483*** | - | - | - | - | - | - | - | - |
| Urban | -0.172 | -0.132 | - | - | - | - | - | - | - |
| Wetlands | 0.066 | -0.002 | -0.364 | - | - | - | - | - | - |
| Budapest | -0.124 | -0.107 | **0.496*** | -0.339 | - | - | - | - | - |
| Age | 0.084 | 0.093 | 0.119 | -0.036 | 0.051 | - | - | - | - |
| Fish | 0.240 | 0.268 | 0.031 | -0.019 | -0.014 | 0.068 | - | - | - |
| AqVeg | 0.088 | 0.060 | 0.061 | -0.010 | 0.053 | -0.070 | 0.246 | - | - |
| ShoreVeg | 0.091 | 0.037 | -0.027 | 0.029 | 0.009 | -0.064 | 0.140 | 0.156 | - |
| Concrete | -0.102 | -0.063 | -0.031 | 0.002 | 0.008 | 0.089 | -0.033 | -0.063 | -0.115 |
| Rubber | 0.386 | 0.348 | -0.042 | -0.036 | -0.042 | 0.007 | 0.243 | 0.143 | 0.168 |
| Plastic | **-0.450*** | -0.377 | 0.097 | -0.032 | 0.082 | -0.067 | -0.217 | -0.090 | -0.084 |
| Metal | -0.108 | -0.094 | 0.010 | 0.035 | 0.028 | -0.036 | -0.090 | -0.066 | -0.031 |
| Natural | 0.091 | 0.026 | -0.090 | 0.117 | -0.064 | 0.029 | -0.123 | -0.186 | -0.069 |
| Draining | -0.121 | -0.080 | 0.028 | 0.040 | 0.026 | 0.185 | 0.076 | -0.024 | 0.011 |

| | Area | Depth | Urban | Wetlands | Budapest | Age | Fish | AqVeg | ShoreVeg |
|---|---|---|---|---|---|---|---|---|---|
| Cleaning | -0.017 | -0.012 | 0.028 | 0.059 | 0.008 | 0.174 | 0.086 | -0.034 | 0.064 |
| Leaves | -0.030 | 0.031 | 0.083 | 0.009 | 0.047 | 0.001 | 0.124 | 0.108 | 0.102 |
| Pruning | 0.013 | -0.026 | -0.002 | -0.017 | -0.017 | 0.068 | -0.149 | 0.016 | -0.019 |
| Circulation | 0.076 | 0.124 | 0.014 | 0.012 | 0.004 | -0.106 | 0.371 | 0.153 | 0.125 |
| Algaecide | -0.043 | -0.052 | 0.029 | 0.006 | 0.017 | -0.117 | 0.174 | 0.062 | 0.042 |

Table S2. (cont.)

| | Concrete | Rubber | Plastic | Metal | Natural | Draining | Cleaning | Leaves | Pruning | Circulation |
|---|---|---|---|---|---|---|---|---|---|---|
| Rubber | -0.270 | - | - | - | - | - | - | - | - | - |
| Plastic | -0.050 | **-0.709*** | - | - | - | - | - | - | - | - |
| Metal | -0.018 | -0.147 | -0.022 | - | - | - | - | - | - | - |
| Natural | -0.040 | -0.332 | -0.050 | -0.009 | - | - | - | - | - | - |
| Draining | 0.030 | -0.007 | 0.062 | 0.043 | -0.103 | - | - | - | - | - |
| Cleaning | 0.029 | 0.001 | -0.018 | 0.055 | -0.031 | 0.396 | - | - | - | - |

| | Concrete | Rubber | Plastic | Metal | Natural | Draining | Cleaning | Leaves | Pruning | Circulation |
|---|---|---|---|---|---|---|---|---|---|---|
| Leaves | 0.043 | 0.089 | -0.027 | -0.007 | -0.164 | 0.074 | 0.079 | - | - | - |
| Pruning | 0.018 | 0.036 | -0.065 | -0.018 | 0.070 | -0.106 | -0.088 | -0.028 | - | - |
| Circulation | -0.018 | 0.110 | -0.053 | -0.014 | -0.120 | 0.144 | 0.125 | 0.213 | -0.165 | - |
| Algaecide | 0.030 | 0.031 | 0.013 | -0.003 | -0.087 | 0.112 | 0.115 | 0.102 | -0.168 | 0.305 |

*strong correlation ($r_s \geq 0.5$)

Area = logarithmic transformation of pond area; Depth = logarithmic transformation of pond water depth; Urban = proportion of urban land cover within a 1-km radius of a garden pond; Wetlands = proportion of wetlands and standing waterbodies within a 1-km radius of a garden pond; Budapest = ponds located within Budapest (1) or elsewhere (0); Age = years since 2021 when pond was built; Fish = deliberate introduction of fish; AqVeg = submerged or floating aquatic vegetation planted within the pond (e.g. water lilies, pondweed); ShoreVeg = vegetation planted within the pond around the shoreline (e.g. sedges, irises); Concrete = pond constructed of concrete; Rubber = pond constructed with rubber (PVC) lining; Plastic = pre-fabricated plastic pond; Metal = pre-fabricated metal pond; Natural = pond with earthen or clay lining; Draining = pond drained at least once by owners; Cleaning = pond bed cleaned (i.e. sediment removed) at least once by owners; Leaves = leaves removed from pond by owners; Pruning = vegetation maintenance (cutting the plants, manually removing plants and algae); Circulation = water circulation device installed in pond (e.g. filtration system, fountain); Algaecide = application of chemical or probiotic algaecides.

**Fig. S1.** Moran's Eigenvector Maps (MEM) for significant eigenvectors: (a) MEM1; (b) MEM3. The cluster at the centre of the maps largely corresponds with the Hungarian capital city, Budapest.

(a)

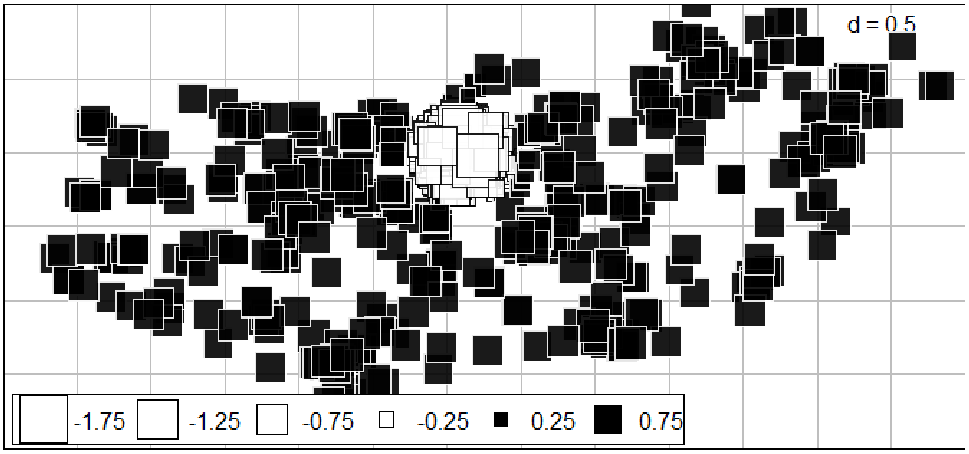


(b)

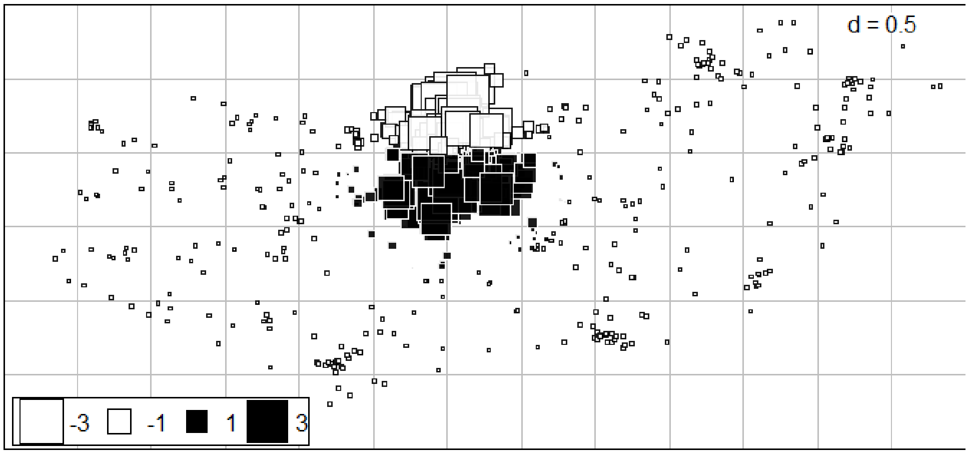

**Fig. S2.** A typical garden pond in Hungary. Note the size, PVC rubber lining, water fountain, aquatic vegetation and ornamental fish.

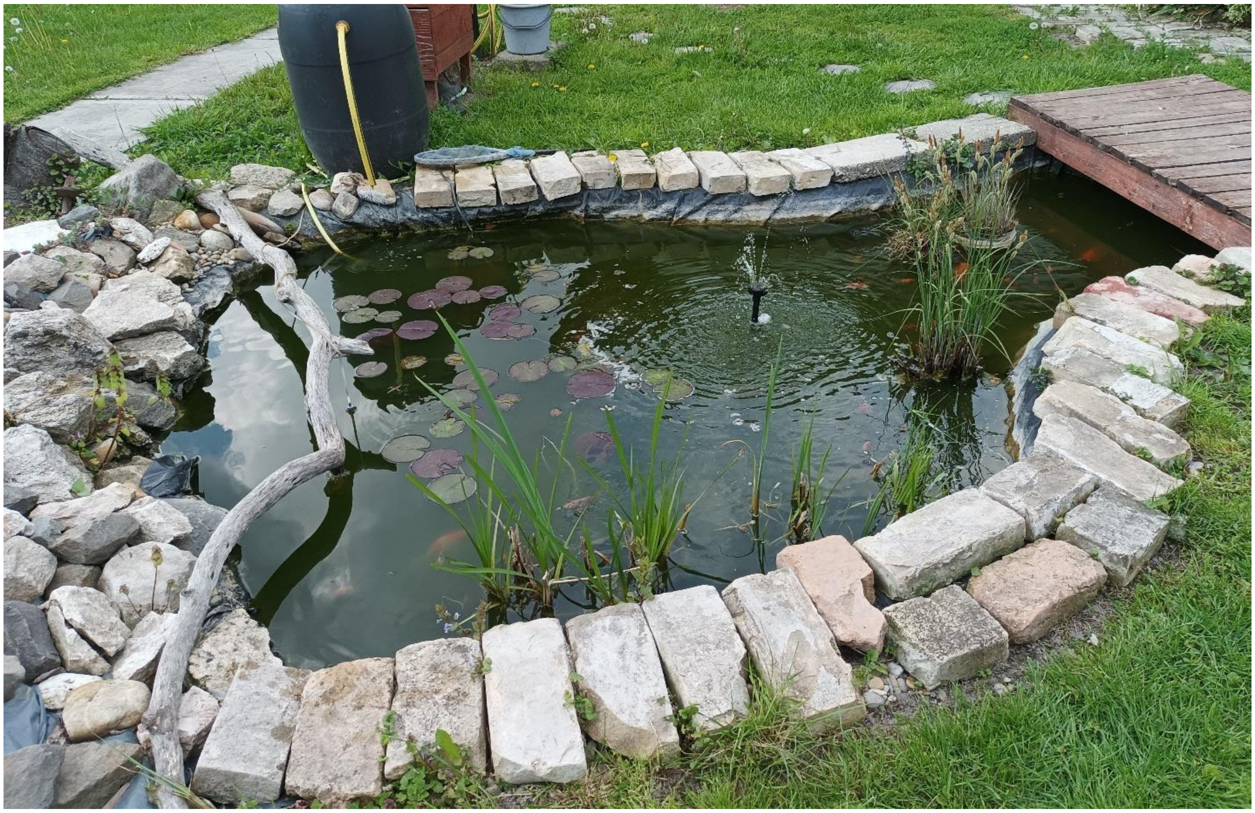
